# Insulin-Like Growth Factor I Modulates Vulnerability to Stress Through Orexin Neurons

**DOI:** 10.1101/2020.02.10.941377

**Authors:** ME Fernandez de Sevilla, J Pignatelli, P. Mendez, J Zegarra-Valdivia, I Torres Alemán

## Abstract

Knowledge of mechanisms involved in vulnerability/resilience to stress disorders is crucial for prevention and treatment schemes. We previously documented that insulin-like growth factor I (IGF-I) is associated to vulnerability to stress both in mice and humans. Since hypothalamic orexin neurons express IGF-I receptors and are involved in responses to stress, we analyzed their role in the modulatory actions of IGF-I on stress. Anxiolytic actions of IGF-I after exposure to a predator were absent in mice lacking IGF-I receptors in orexin neurons (Firoc mice). Based on these observations we speculated that Firoc mice may be prone to develop fear-related disturbances, including post-traumatic stress disorder (PTSD)-like symptoms when confronted to fear learning, a process that is postulated to be altered in PTSD. Firoc mice submitted to fear conditioning showed increased freezing responses, suggesting aberrant fear learning. Exaggerated freezing was accompanied by increased levels of orexin, together with enhanced c-fos staining of these neurons –an indicator of increased cell activity, and of noradrenergic neurons of the locus coeruleus nucleus, a region downstream of orexinergic activation. After fear conditioning, Firoc mice developed PTSD-like behavioral traits such as prolonged context-dependent fear and post-stress anhedonia. Since abnormal fear learning was ameliorated by chemogenetic (DREADD) inhibition of orexin neurons, reduced IGF-I input to orexin neurons in Firoc mice seems to enhance their excitability to fear-related inputs. Collectively, these results suggest that IGF-I input to orexin neurons is an important determinant of vulnerability to stress disorders, which provides additional targets for therapy of these high social impact conditions.

## Introduction

Exposure to a traumatic event affects a large part of the world population, but only a relatively small proportion will develop a post-traumatic stress disorder (Shalev et al., 2017). However, specific subsets of the population are at greater risk, such as soldiers exposed to combat (Yurgil et al., 2014). Unfortunately, treatment of this mental condition remains largely unsatisfactory (DePierro et al., 2019). Mechanisms of vulnerability and resilience to PTSD are therefore crucial to treat, and even prevent, this psychiatric illness. Since we recently found that IGF-I is associated to vulnerability to stress in mice and humans (Santi et al., 2018b), we have explored underlying mechanisms with the hope to gain insight into novel processes affecting vulnerability/resilience to stress in general, and to PTSD development more specifically.

Modulatory actions of IGF-I on mood are increasingly recognized (Lin et al., 2014; van Varsseveld et al., 2015; Kondo et al., 2017; Santi et al., 2018b). The relation of circulating IGF-I with mood encompasses multiple angles, such as anxiolysis (Baldini et al., 2013; Santi et al., 2018b), arousal (Bellar et al., 2011), playfulness (Burgdorf et al., 2010), or depression (Deuschle et al., 1997; Kondo et al., 2017). The latter aspect is profusely documented in the literature. In search for possible cellular targets of IGF-I in mood modulation we focused on the hypothalamus, a brain area that expresses abundant IGF-I receptors (http://mouse.brain-map.org/experiment/show?id=69735263) and is central in neuroendocrine regulation of stress (Ulrich-Lai and Herman, 2009). Within the hypothalamus, orexin neurons are involved in responses to stress (Ji et al., 2019) and pharmacological modulation of their activity has been proposed as possible therapy for PTSD (Flores et al., 2015; Soya and Sakurai, 2018; Cohen et al., 2020). These neurons are also involved in mood regulation in general (Harris et al., 2005; Johnson et al., 2010; Blouin et al., 2013), and mood-related traits such as motivated behaviors (Mahler et al., 2014; Sakurai, 2014), or arousal (Yamanaka et al., 2003). Recently, orexin neurons have been shown to be directly involved in PTSD-like responses in rats (Cohen et al., 2020). Thus, we hypothesized that orexin neurons may participate in the regulation of stress responses by IGF-I.

In this work we report that a functional link between IGF-I and orexin is involved in vulnerability to stress. Mice with disrupted IGF-IR activity in orexin neurons developed PTSD-like traits including exacerbated fear conditioning, delayed extinction of fear, and post-stress anhedonia. Abnormally high orexin activation underlies this phenotype, as their chemogenetic inhibition ameliorated it.

## Results

### Anxiolytic actions of IGF-I involve orexin neurons

We recently found that IGF-I directly modulates the activity of orexin neurons (submitted). Since IGF-I reduces anxiety in mice exposed to a predator (Santi et al., 2018b), a natural anxiogenic stimulus (Blanchard et al., 2003), we administered IGF-I (icv) to mice expressing an inactive IGF-IR in orexin neurons (Firoc mice) and their control littermates, before exposing them to a rat (Figure 1). We observed reduced anxiety in littermate controls treated with IGF-I, but not in Firoc mice, as determined by reduced entrance to the open arms in the elevated plus maze (EPM; Figure 1C). Other parameters measuring fear behavior during predator exposure (freezing and grid contacts), and a subsequent test in the open field indicating enhanced anxiety, were also significantly ameliorated by IGF-I treatment in control, but not in mutant mice (Figure 1B,D). Lack of anxiolytic actions of IGF-I in Firoc mice suggests that these neurons are involved in stress regulation by IGF-I.

**Figure 1.**
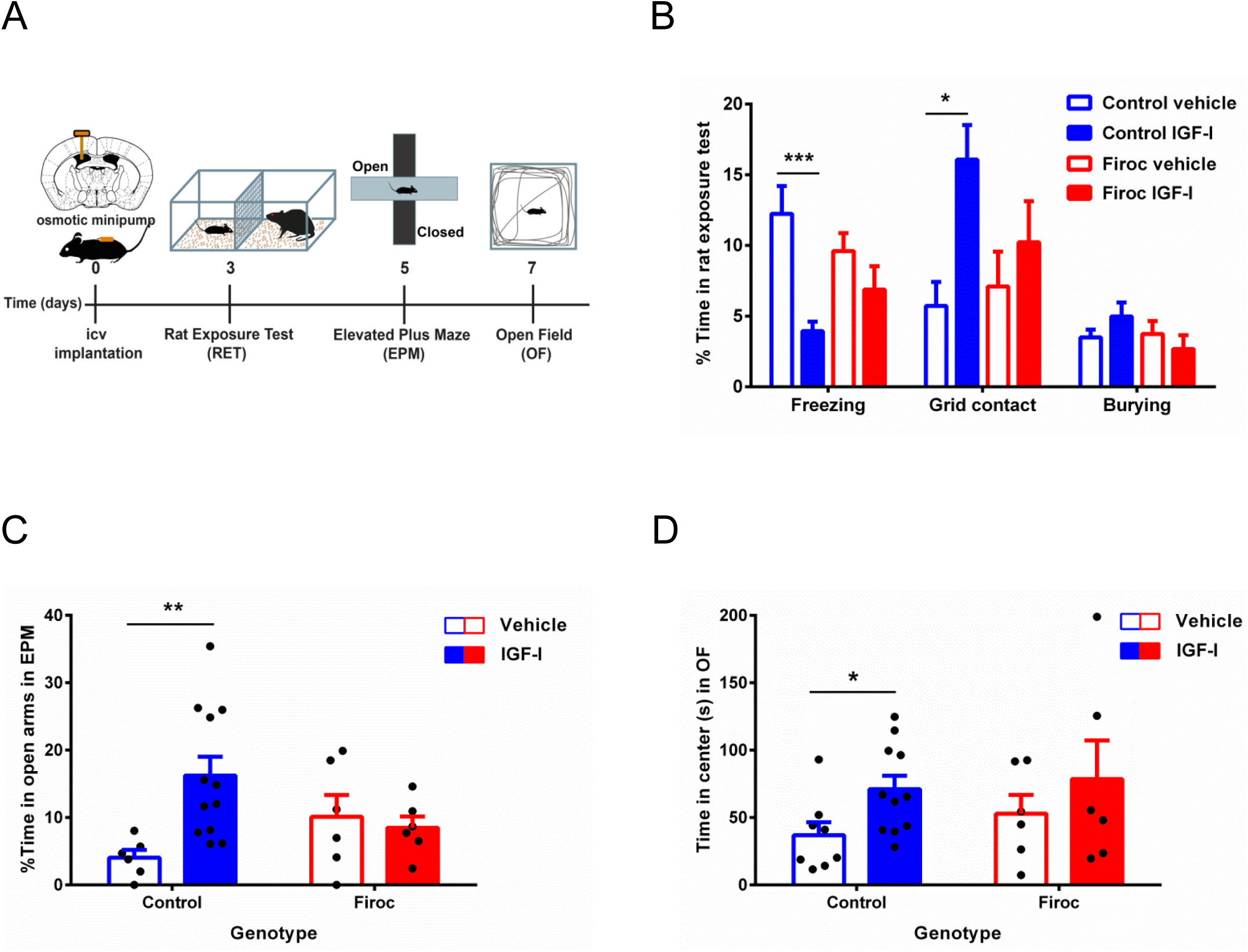
Anxiolytic actions of IGF involve orexin neurons. **A**, Time-line of experiments. **B**, Ethogram obtained during the rat exposure test (RET). Intracerebroventricular administration of IGF-I attenuated fear behaviors in control animals but not in Firoc mice; freezing was significantly decreased and grid contacts increased. Burying behavior was not altered by predator exposure. **C**, Anxiety levels were measured 2 days after RET in the EPM test as percentage of time spent in the open arms. Only IGF-I-treated control animals spent more time in the open arms. **D**, Similarly, less anxiety in the OF test was observed 4 days after predator exposure only in control animals treated with IGF-I; i.e.: greater amount of time spent in the center of the arena (t-test; *p<0.05; **p<0.01; ***p<0.001 in this and following figures).

### IGF-I modulates orexin responses to fear learning

Based on the above observations, we hypothesized that Firoc mice could show disturbed fear learning. Hence, we submitted them to fear conditioning (Figure 2A), and found that Firoc mice showed increased fear responses, as measured by time spent in freezing behavior (Figure 2B and Suppl Fig 1A,B). Firoc mice exposed to fear conditioning also showed increased number of activated orexin (double labeled c-fos/orexin neurons; Figure 3A-C), and locus coeruleus (LC) neurons (double labeled c-fos/TH cells, Figure D-F), a prominent downstream connection of orexin neurons in fear responses (Soya et al., 2017).

**Figure 2.**
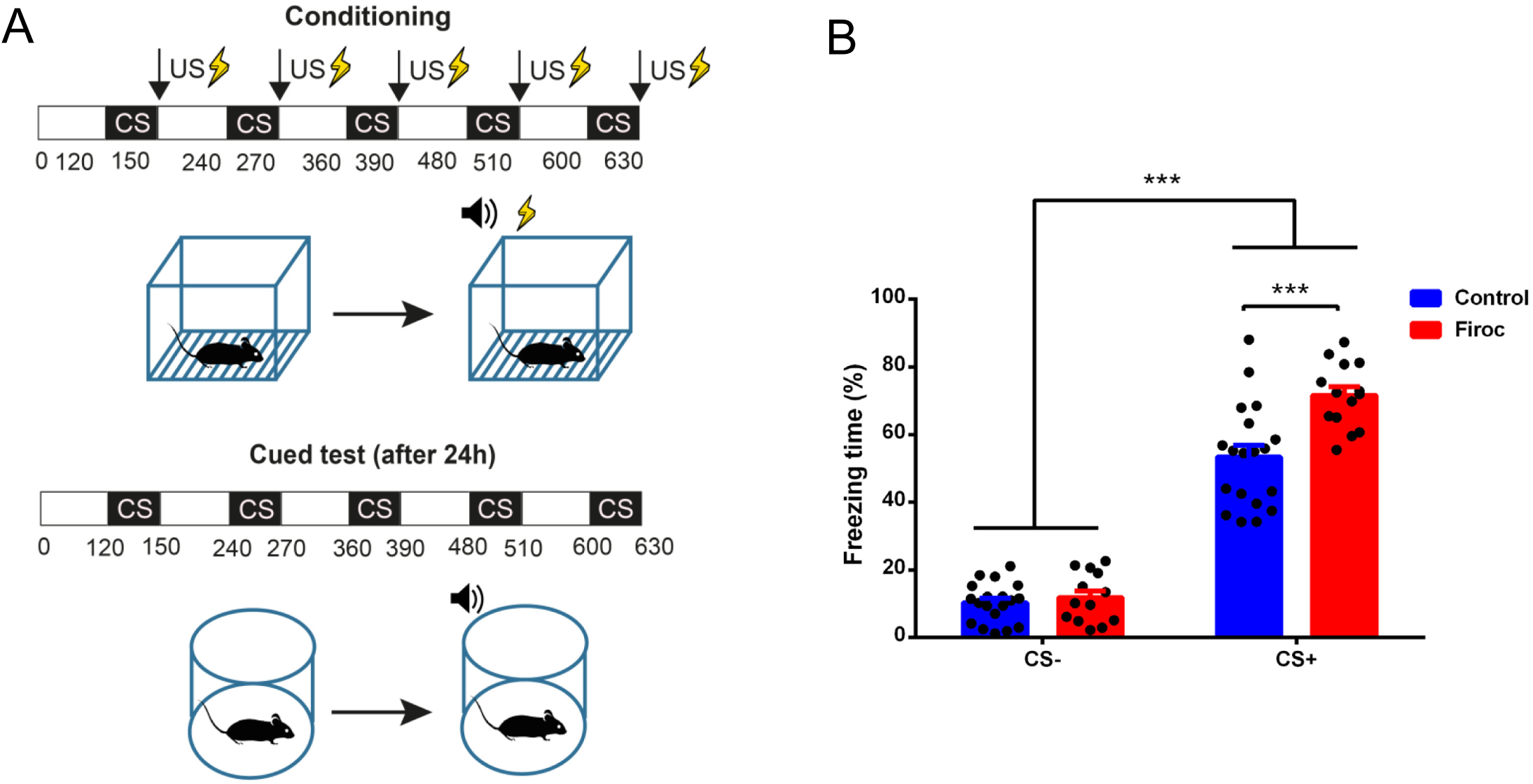
Modulation by IGF-I of fear conditioning through orexin neurons. **A**, Schematic drawing of the protocol used in cued fear-conditioning. Top: A neutral conditioned stimulus (CS), a tone of 80 dB, is presented together with an aversive unconditioned stimulus (US), an electrical footshock of 0.3 mA. Bottom: as a consequence of this US–CS association, a new exposure to the CS in the absence of the US in a different context elicits a conditioned freezing response. **B**, Firoc mice presented a higher percentage of freezing time in the cued test phase during CS periods (see Supplementary Figure 1A,B for additional information; n=14-19 mice/group; Two-way-ANOVA followed by Tukey’s multiple comparisons test).

**Figure 3.**
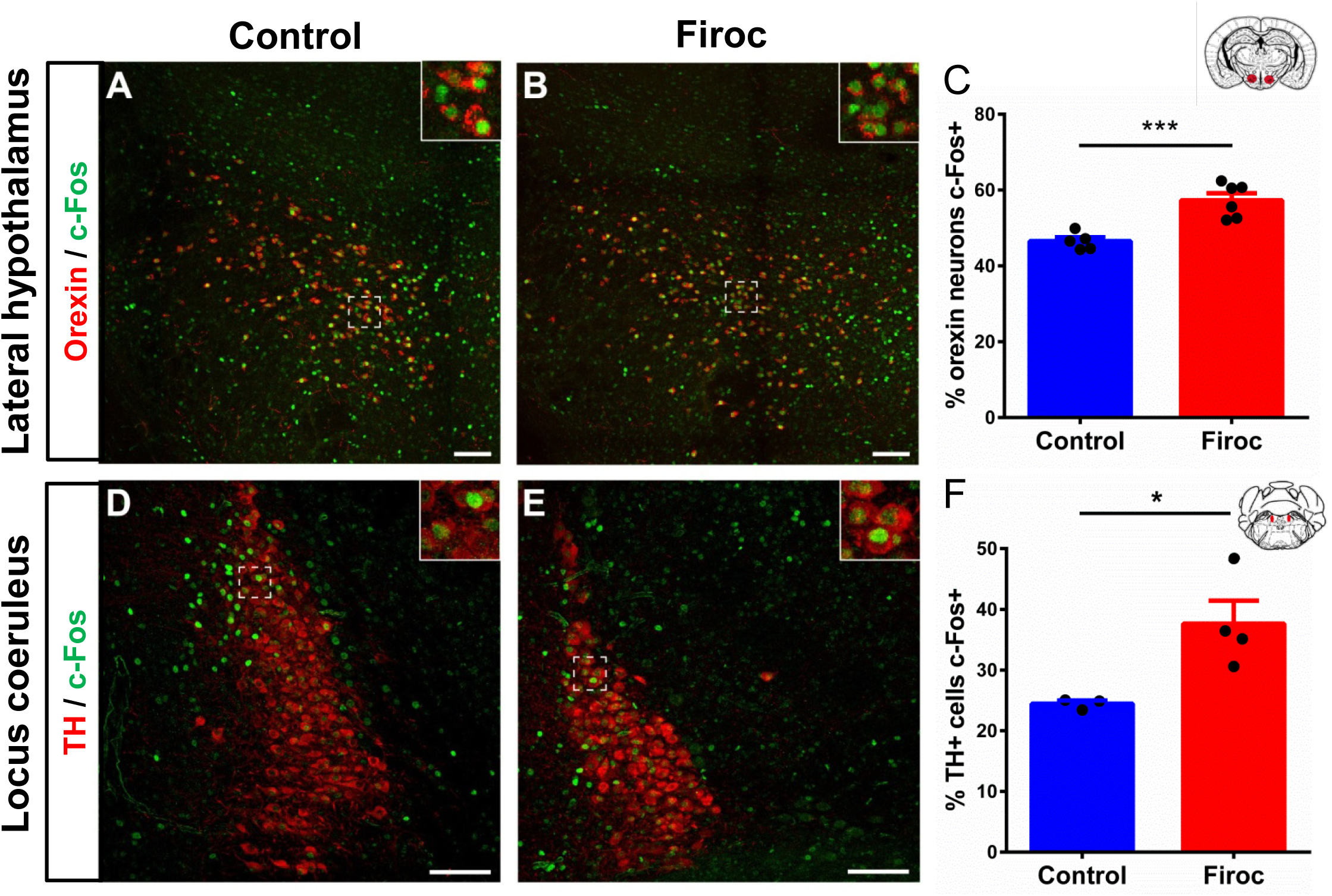
Fear-induced activation of orexin and LC neurons is modulated by IGF-I through orexin neurons. **A, B**, Double-stained c-fos (green) and orexin cells (red) in the lateral hypothalamus of control (A) and Firoc mice (B) exposed to fear conditioning. **C**, Number of c-fos^+^ /orexin^+^ cells (expressed as percent of total orexin^+^ cells) was significantly increased in Firoc mice after fear conditioning (n=5-6). **D**,**E**, Double-stained c-fos (green) and TH cells (red) in the LC of control (D) and Firoc mice (E) exposed to fear conditioning. **F**, Number of c-fos^+^ /TH^+^ cells (expressed as percent of total TH^+^ cells) was significantly increased in Firoc mice after fear conditioning. (n= 3-4). Scale bar: 100μm.

Since these observations suggested increased activation of orexin neurons in Firoc mice submitted to fear conditioning, we chemogenetically inhibited orexin activity. After bilateral injection of an inhibitory DREADD-Cre virus into the lateral hypothalamus of Orexin-Cre (controls) and Firoc mice, we administered CNO to fear-conditioned mice and found that Firoc mice normalized their fear responses (Figure 4A). Control littermates and Firoc mice injected with mCherry viruses show normal or exaggerated fear responses, respectively, as expected. These data suggest that orexin neurons in Firoc mice are overactivated.

**Figure 4.**
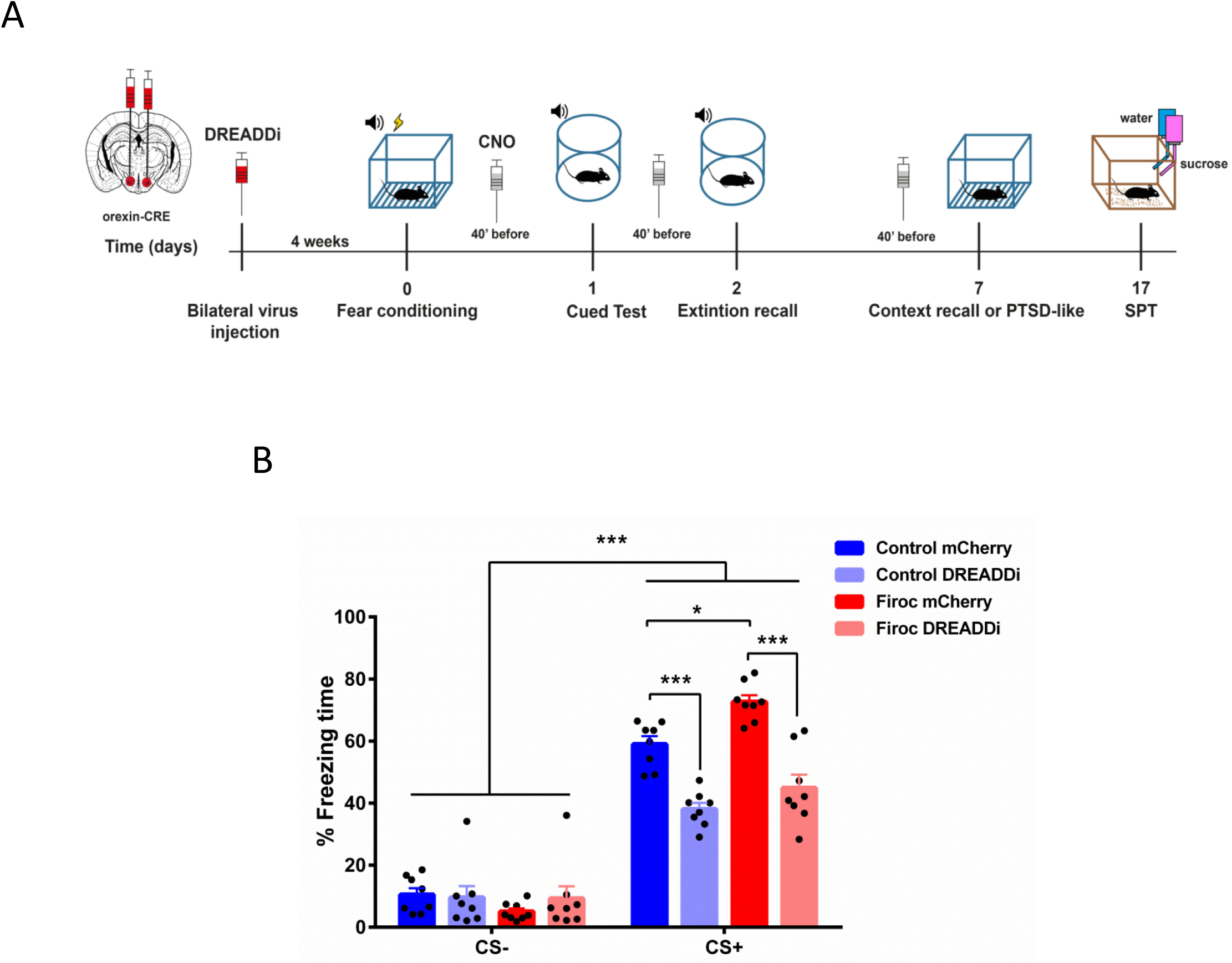
IGF-I modulates excitability of orexin neurons. **A**, Time-line of experimental procedures followed in DREADD experiments in this and the following figure. **B**, Chemogenetic inhibition of orexin neurons in Firoc mice attenuates abnormal freezing responses. Control littermates and Firoc mice injected with mCherry control virus show normal and increased fear learning, respectively (n=7-8; Two-way-ANOVA followed by Tukey’s multiple comparisons test).

### Firoc mice display PTSD-like traits

Abnormal fear learning is suggested to underlie PTSD (Shalev et al., 2017). Taking advantage that chemogenetic inactivation of orexin neurons in Firoc mice rescues exaggerated fear learning, we examined possible development of PTSD-like traits and its attenuation with DREADD inhibition in Firoc mice. Extinction of context-dependent fear responses (context recall) 1 week later were markedly attenuated in Firoc mice injected with mCherry virus as compared to mCherry-injected littermate controls, indicating abnormal retention of learned fear (Figure 5A), a trait seen in PTSD patients. Interestingly, these mice also showed higher levels of orexin in the hypothalamus after fear conditioning (Figure 5B). We also found that after fear conditioning, Firoc mice developed anhedonia, as measured by the sucrose-preference test (Figure 5C), another characteristic of PTSD patients. Importantly, Firoc mice did not show anhedonia under basal conditions (not shown).

**Figure 5.**
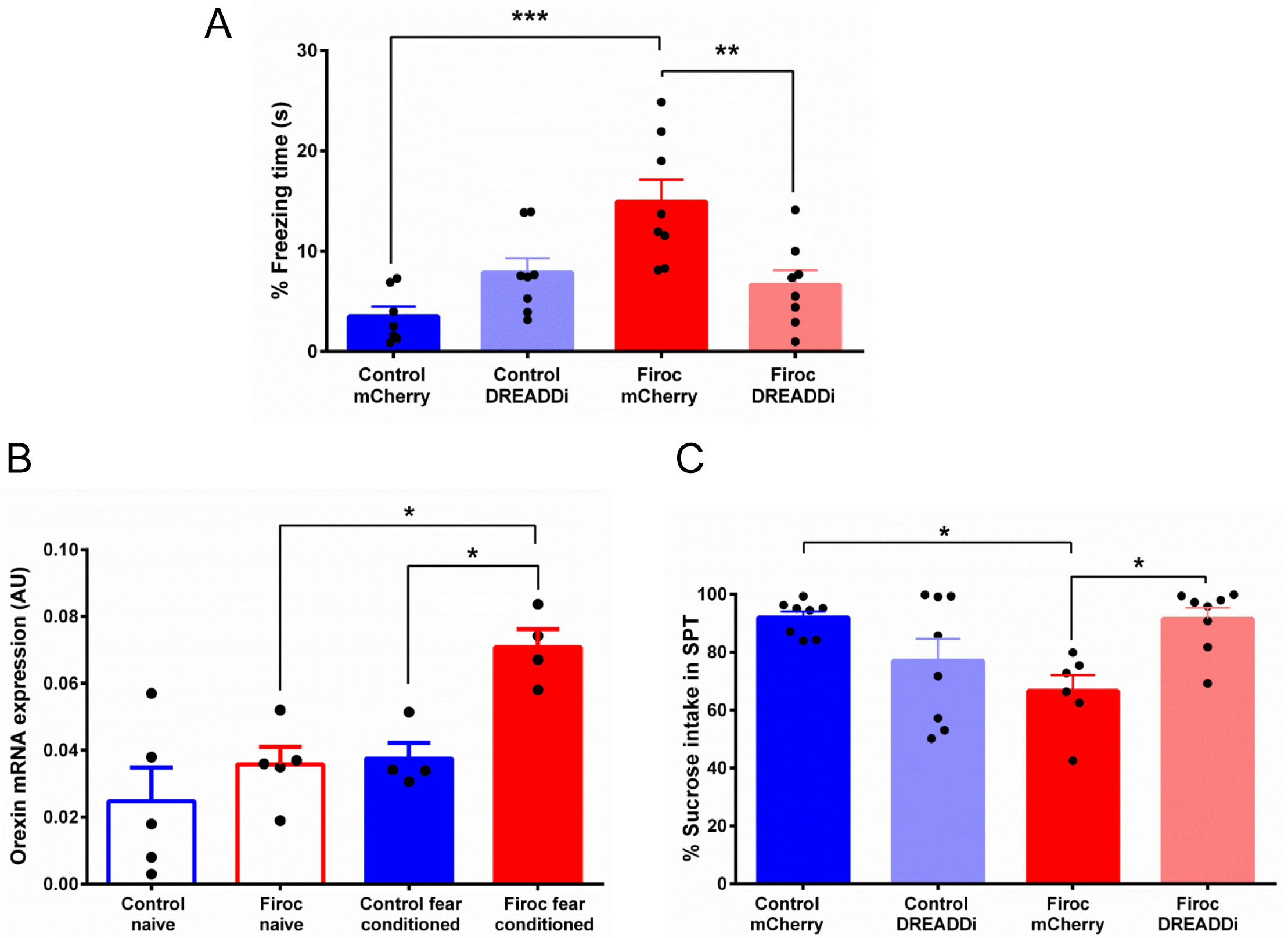
Firoc mice show PTSD-like traits. **A**, One week after fear conditioning, animals were retested to determine extinction recall by placing them in the same context. Firoc mice injected with mCherry show reduced extinction that was normalized after DREADD inhibition (n=7-8; One-way-ANOVA followed by Tukey’s multiple comparisons test). B, Firoc mice submitted to fear conditioning show increased orexin expression in the hypothalamus (n= 5-6; Mann Whitney U Test). **C**, Firoc mice showed anhedonia as tested in the SPT that was rescued with DREADD-mediated chemogenetic inhibition of orexin neurons (n=6-8; One-way-ANOVA followed by Tukey’s multiple comparisons test).

## Discussion

These results indicate that IGF-I modulates responses to stress through hypothalamic orexin neurons. Regulation of orexin activity by IGF-I probably forms part of homeostatic regulation of these neurons (Azeez et al., 2018). We show that the IGF-I receptor in orexin neurons modulate both acquired (fear learning), and innate fear (predator exposure) responses, providing a specific molecular mechanism for vulnerability to aberrant responses to stress. Indeed, blunted IGF-I input to orexin neurons in Firoc mice favors the appearance of PTSD-like behavior, including retention of fear memories, anhedonia and altered sensitivity to dexametasone, three traits observed in human patients (Pietrzak et al., 2014). Significantly, the PTSD-like phenotype of Firoc mice is normalized by chemogenetic inhibition of orexin over-activity. These findings reinforce the previously reported use of orexin receptor antagonists for treatment of this important mental illness (Flores et al., 2015; Soya and Sakurai, 2018).

Conversely, recent observations pose orexin activation in response to fear stimuli (predator smell) as a resilient mechanism (Cohen et al., 2020). These two apparently opposing observations may be reconciled with our findings. Since anxiolytic actions of IGF-I require signaling onto orexin neurons (Figure 1), IGF-I may modulate orexin activity in response to fear stimuli to cope with them, as suggested by Cohen et al (Cohen et al., 2020). In turn, in the absence of IGF-I input (Firoc mice), orexin activity becomes maladaptive; i.e.: greater expression of orexin and increased number of double-labelled c-fos^+^/orexin^+^ and c-fos^+^/TH^+^ cells are found in Firoc mice after fear conditioning, as compared to littermates. Moreover, Firoc mice show impaired fear extinction. Thus, inappropriate activation of the orexin-LC circuitry will lead to aberrant fear behavior, as suggested by others (Soya et al., 2017).

The fact that a pleiotropic neurotrophic factor such as IGF-I (Fernandez and Torres-Aleman, 2012) modulates the activity of orexin neurons, considered also a multitasking system (Sakurai, 2014), help to explain the diversity of actions of IGF-I in the brain, including regulation of mood. Indeed, numerous reports relate IGF-I with affective disorders (Deuschle et al., 1997; Hoshaw et al., 2008; Kim et al., 2013; Lin et al., 2014; Bot et al., 2016; Chigogora et al., 2016), and specific potential mechanisms have been started to unfold. For example, recent observations indicate that IGF-I participates in depression through its role in the serotonin system (Kondo et al., 2017), while others related it to its involvement in adult hippocampal neurogenesis (Anderson et al., 2002). Thus, our observations widen the number of potential mechanisms whereby IGF-I intervenes in mood homeostasis, now including the orexinergic system, which emphasizes the complexity of mechanisms mediating the actions of this growth factor, even within the same category of activities.

## Materials and Methods

### Materials

Antibodies used include rabbit polyclonal anti-c-Fos (1:500 dilution, sc-52, Santa Cruz Biotechnology), monoclonal anti-orexin-A (1:100 dilution, sc-80263, Santa Cruz), and polyclonal anti-tyrosine hydroxylase (1:1000 dilution, MAB318, Millipore). Human recombinant IGF-I was from Pre-Protech (USA). For hdM4Di (DREADD) experiments, clozapine-N-oxide (CNO) from Tocris at 2mg/kg dissolved in saline 0.9% was injected intraperitoneally 40 min before test sessions 24h and 1week after fear conditioning.

### Animals

Adult female and male C57BL/6J mice (Harlan Laboratories, Spain), and Cre/Lox mice lacking IGF-I receptors in orexin neurons (Firoc mice) were used. Firoc mice were obtained by crossing Orexin-Cre mice (a kind gift of Dr T Sakurai, Tsukuba Univ, Japan; (Matsuki et al., 2009) with IGF-IR^f/f^ mice (B6, 129 background; Jackson Labs; stock number: 012251) as explained in detail elsewhere (submitted). Orexin neurons in Firoc mice do not respond to systemic IGF-I administration, as assessed by c-fos and phospho-Akt expression (not shown).

Genotyping of Firoc mice was performed using 5’-GGTTCGTTCACTCATGGAA AATAG-3’, and 5’-GGTATCTCTGACCAGAGTCATCCT-3’ for Orexin-Cre and 5’-CTTCCCAGCTTGCTACTCTAGG-3’, and 5’-CAGGCTTGCAATGAGACATGGG-3’ for IGF-IR^f/f^. DNA from brain tissue was isolated using Trizol Reagent and ethanol precipitation. 10ng of genomic DNA was used in a PCR reaction containing 1X reaction buffer, 1µM of each primer, 0.2mM of dNTPS and 0.75ul of DFS-Taq DNA polymerase (Bioron, GmbH). The thermocycler program was 92°C, 3 min and 30 cycles of 94°C, 30sec; 65°C, 30sec; 72°C, 30sec, after that a final extension step at 72°C for 2 min was performed. Amplicons were analyzed in 3% agarose gels stained with SYBRsafe (Thermofisher, Inc).

Animals were housed in species-specific standard cages (mice 5 per cage; rats 1–2 rats per cage), and kept in a room with controlled temperature (22°C) under a 12-12h light-dark cycle. All animals were fed with a pellet rodent diet and water *ad libitum*. All experimental protocols were performed during the light cycle. Mice were handled for 3 days prior to any experimental manipulations. Animal procedures followed European guidelines (86**/**609**/**EEC and 2003**/**65**/**EC, European Council Directives) and were approved by the local Bioethics Committee (Government of the Community of Madrid).

### Viral injection

For chemogenetic experiments using DREADD, a viral construct (pAAV-hSyn-DIO-hM4D(Gi)-mCherry; AAV5; 8.6 × 10^12^ viral infective units/ml) was locally injected bilaterally to inactivate orexin-cre neurons in Orexin-Cre (littermates) and transgenic mice (Firoc). As control viruses, we used pAAV-hSyn-DIO-mCherry (AAV5). Both viral constructions were obtained from Addgene. CNO efficacy in orexin neurons was confirmed in acute slices obtained from injected mice. Slices for electrophysiological recordings were prepared from 2-months old Orexin-Cre mice, 4 weeks after injection of the DREADD-mCherry virus into the lateral hypothalamus. Brains were quickly removed and coronal slices (250 µm) containing the lateral hypothalamus were cut with a vibratome (4°C) in a solution containing: 234 mM sucrose, 11 mM glucose, 26 mM NaHCO3, 2.5 mM KCl, 1.25 mM NaH2PO4, 10 mM MgSO4, and mM 0.5 CaCl2 (equilibrated with 95% O2–5% CO2). Recordings were obtained at 30-32 °C from orexin^+^ neurons identified using fluorescence microscopy (mCherry+) in oxygenated artificial cerebrospinal fluid containing the following: 126 mM NaCl, 26 mM NaHCO3, 2.5 mM KCl, 1.25 mM NaH2PO4, 2 mM MgSO4, 2 mM CaCl2 and 10 mM glucose (pH 7.4). Patch-clamp electrodes contained intracellular solution composed of: 131 mM K gluconate, 5 mM KCl, 4 mM MgCl2, 10 mM HEPES, 4 mM EGTA, 2 mM MgATP, and 0.3 mM Na2GTP (pH 7.3) corrected with KOH (290 mOsm). Positive and negative currents were injected during 600 ms to calculate action potential frequency and input resistance. Clozapine N-Oxide (2µM) was applied through perfusate disolved in ACSF. Signals were amplified, using a Multiclamp 200B patch-clamp amplifier (Axon Instruments, Foster City, California, United States), sampled at 20 kHz, filtered at 10 kHz, and stored on a PC. Data were analyzed using pClamp (Axon Instruments). We confirmed CNO-mediated inhibition of orexin neurons in hypothalamic slices of Orexin-Cre mice transduced with pAAV-hSyn-DIO-hM4D(Gi)-mCherry (Suppl Fig 1 C-F).

### Surgery

For all surgeries, mice were anaesthetized with isoflurane (Zoetis) administered with a nose mask (David Kopf Instruments, France), and placed on a stereotaxic frame (Stoelting Co) on a heating pad and tape in their eyes to protect them from light. For viral expression, mice were injected with a 5 µl Hamilton syringe bilaterally into the orexin nuclei (AP= −1.4; ML= ± 0.9; DV= −5.4) 4 weeks before experiments. For DREADD viral expression, 300 nl were bilaterally infused at a rate of 100 nl/min and the Hamilton syringe withdrawn 10 min later.

## Behavioral tests

### Open field

Locomotion and exploratory behavior were evaluated placing the animal in an open field arena (42 cm× 42 cm × 30 cm, Versamax; AccuScan Instruments, Inc.) for 10 min. All parameters were quantified automatically with the software provided.

### Cued-fear conditionin

Experiments were performed at the same time of the day during the light phase with 3-4 months-old male and female mice (n=14-19 mice/genotype, balanced sex). The training and test sessions were video recorded. Mice were placed in a shuttle box chamber (AccuScan Instruments) for 120 sec before exposing them to a neutral conditioned stimulus (CS+) -a tone of 80 dB for 30s, presented together with an aversive unconditioned stimulus (US) -an electrical footshock of 0.3 mA/2 sec. Training consisted of 5 consecutive trials with 90 sec intertrials (CS-). Animals remained in the chamber for 5 min more before returning them to their home cages. Twenty four hours later, fear conditioning was tested placing the animals in a different context using the same protocol but without the unconditioned stimulus (US). As a consequence of the US–CS association, mice displayed freezing behavior, which was scored for every trial (CS+), and intertrial (CS-). Freezing behavior was defined by the absence of movement except of breathing and heart beating. Context-dependent extinction was tested 1 week later (delayed extinction). Mice were placed in the same context for 5 min to measure their freezing behavior. The latter unveils PTSD-like behaviour when it is abnormally retained.

### Predator exposure test

Control littermates and Firoc mice implanted with osmotic minipumps (either with saline or IGF-I, see below), were introduced into a box for 10 min with two compartments separated by a plastic grid, one containing a rat (predator) and the other empty. The box contained bedding of the rat with urine and feces. All sessions were recorded, and every rat exposed to a maximum of 4 mice/day. Grid contacts, freezing time and bedding burying behaviors of exposed mice were scored.

### Elevated plus maze

Two days after exposure to the rat, mice were introduced in a maze of 40 cm from the floor with two opposing protected (closed) arms of 30 cm (length) × 5 cm (wide) × 15.25 (height), and two opposing unprotected (open) arms of 30 cm (length) × 5 cm (wide). Each animal was introduced in the center of the maze for 10 min. Stress was scored as time animals spent in closed arms. All measures were recorded with an automated video-tracking system (Video Tracking Plus Maze Mouse; Med Associates, USA).

### Sucrose preference test

Mice were given 2 bottles of water for 3 days, and then 2 bottles of 2% sucrose for 2 days. Afterwards, mice were deprived of food and water for 18h and then, they were presented a bottle of water and a bottle of 2% sucrose during 2h. The position of the bottles was switched after 1h. Bottles were placed in the active phase. Water and sucrose consumption was recorded and sucrose preference was defined as the ratio of the weight of sucrose intake to the weight of total intake of liquid (water and sucrose).

### Immunocytochemistry

Mice were perfused transcardially under deep anesthesia (sodium pentobarbital, 50 mg/kg, i.p.) with 100 ml of saline buffer 0,9% followed by 100 ml of 4% paraformaldehyde (PFA) in 0.1N, pH 7.4 phosphate buffer (PB), 90 min after the cued-fear conditioning test. This time was selected as optimal to see c-Fos staining. The brains were removed, postfixed overnight at 4°C in the same fixative, and cut at 50 µm thick sections on a vibratome (Leica VT 1000S). Sections were kept at 4°C, immersed in 0.1N PB with 0.02% sodium azide, until processing. Serial coronal free-floating sections were rinsed in 0.1N PB for 10 min, and a blocking solution containing 10% normal donkey serum (NDS) and 0.4% Triton X-100 in 0.1N PB was added, and maintained at room temperature for 2 h. Thereafter, sections were incubated overnight at 4°C in the same solution with the corresponding primary antibodies. The next day, sections were washed 3 times with PB-0.4% Triton X-100 (PB-T), and incubated with secondary antibodies AlexaFluor-488 donkey anti-rabbit (1:1000 dilution, A-21206, Life Technologies), and AlexaFluor-594 donkey anti-mouse (1:1000 dilution, A-21203, Life Technologies) for 2 h at room temperature. After the incubation, slices were washed 3 times with PB-T and incubated 5 min with Hoescht (1:500 dilution, Life Technologies). Finally, sections were washed 3 times with PB and mounted onto glass slides coated with gelatin in Gelvatol mounting medium. Images were taken with confocal microscopy (SP5, Leica Microsystems, Germany) and cell counting was performed with Imaris software.

### qPCR

Animals were anesthetized with pentobarbital (50 mg/kg, ip). After dissecting the brain, the different brain areas were collected and frozen at −80°C until use. Tissue RNA was extracted with Trizol (Life Technologies, USA), as described elsewhere (Santi et al., 2018a). cDNA was synthesized from 1μg of RNA of each sample following the manufacturer’s instructions (Hight Capacity cDNA Reverse Transcriptic Kit; Applied Biosystems). Fast Real-time qPCR was performed using the SYBR Green method (Fast SYBR Green Master Mix, Applied Biosystems) with the QuantStudio 3 Real-Time PCR System (Applied Biosystems). Relative mRNA expression was determined by the 2^−ΔΔCT^ method (Pfaffl, 2001), and normalized to GAPDH levels.

### IGF-I administration

Human recombinant IGF-I (Pre-Protech, USA) or vehicle (saline) were administered intracerebroventricularly (icv, 50 µg/kg/day) using Alzet osmotic mini-pumps for 7 days (Model 1007D), and the brain infusion kit 3 (Alzet, USA) with the following stereotaxic coordinates: AP= −0.5; ML= 1.1; DV= −2.5. Alzet pumps were implanted subcutaneously between the scapulae.

### Statistics

Statistical analyses were performed with GraphPad Prism 6 software (San Diego, USA). Student’s t-test (for comparing two groups) test for comparing two groups or either 1-way or 2-way ANOVAs (for more than 2 groups) followed by Tukey’s multiple comparison test as a post hoc were performed. For non-normally distributed data, we used the Mann Whitney U. All results are shown as mean ± standard error (SEM) and significant values as: *p<0.05; **p<0.01; ***p<0.001.

## Acknowledgements

We are thankful to M. Garcia and R. Cañadas for technical support. This work was funded by a grant from Ciberned, and from SAF2016-76462 (AEI/FEDER; MINECO). JAZ-V acknowledges the financial support of the National Council of Science, Technology and Technological Innovation (CONCYTEC, Perú) through the National Fund for Scientific and Technological Development (FONDECYT, Perú).

## LEGENDS TO FIGURES

**Supplementary Figure. A, B**, Fear-conditioning test: freezing time was scored every inter-trial of 90s (CS-), and trial (CS+) of 30s, both during the acquisition (A), and cued phases (B) (n=14-19 mice/group; *p<0.05; **p<0.01vs controls). **C-F**, CNO activity in orexin neurons. **C**, Membrane potential response to a −25 pA current step of an iDREADD-expressing LH neuron in Orexin-Cre mice previously injected with AAV-AAV5-DIO-iDREADD-mCherry. Traces represent the average of 3 consecutive sweeps before (Control, black) and after 2 µM Clozapine N-Oxide application (CNO, red). **D**, Relative change in input resistance (Rin) in response to CNO application (red arrow). **E**, Firing in response to a depolarizing current step (+15 pA) before (Control, black) and after 2µM CNO application (CNO, red). Traces represent the superposition of 3 consecutive sweeps in each condition. **F**, Relative change in firing frequency in response to CNO application (red arrow). N=4 neurons. Scale bars, 100 ms, 10 mV.

## References

Anderson MF, Aberg MA, Nilsson M, Eriksson PS (2002) Insulin-like growth factor-I and neurogenesis in the adult mammalian brain. Brain Res Dev Brain Res 134:115–122.

Azeez IA, Del Gallo F, Cristino L, Bentivoglio M (2018) Daily Fluctuation of Orexin Neuron Activity and Wiring: The Challenge of “Chronoconnectivity”. Front Pharmacol 9:1061.

Baldini S, Restani L, Baroncelli L, Coltelli M, Franco R, Cenni MC, Maffei L, Berardi N (2013) Enriched early life experiences reduce adult anxiety-like behavior in rats: a role for insulin-like growth factor 1. J Neurosci 33:11715–11723.

Bellar D, Glickman EL, Juvancic-Heltzel J, Gunstad J (2011) Serum insulin like growth factor-1 is associated with working memory, executive function and selective attention in a sample of healthy, fit older adults. Neuroscience 178:133–137.

Blanchard DC, Griebel G, Blanchard RJ (2003) The Mouse Defense Test Battery: pharmacological and behavioral assays for anxiety and panic. Eur J Pharmacol 463:97–116.

Blouin AM, Fried I, Wilson CL, Staba RJ, Behnke EJ, Lam HA, Maidment NT, Karlsson KAE, Lapierre JL, Siegel JM (2013) Human hypocretin and melanin-concentrating hormone levels are linked to emotion and social interaction. Nat Commun 4:1547.

Bot M, Milaneschi Y, Penninx BW, Drent ML (2016) Plasma insulin-like growth factor I levels are higher in depressive and anxiety disorders, but lower in antidepressant medication users. Psychoneuroendocrinology 68:148–155.

Burgdorf J, Kroes RA, Beinfeld MC, Panksepp J, Moskal JR (2010) Uncovering the Molecular Basis of Positive Affect Using Rough-and-Tumble Play in Rats: A Role for Insulin-Like Growth Factor I. Neuroscience 126:769–777.

Cohen S, Matar MA, Vainer E, Zohar J, Kaplan Z, Cohen H (2020) Significance of the orexinergic system in modulating stress-related responses in an animal model of post-traumatic stress disorder. Translational Psychiatry 10:10.

Chigogora S, Zaninotto P, Kivimaki M, Steptoe A, Batty GD (2016) Insulin-like growth factor 1 and risk of depression in older people: the English Longitudinal Study of Ageing. Transl Psychiatry 6:e898.

DePierro J, Lepow L, Feder A, Yehuda R (2019) Translating Molecular and Neuroendocrine Findings in Posttraumatic Stress Disorder and Resilience to Novel Therapies. Biological psychiatry 86:454–463.

Deuschle M, Blum WF, Strasburger CJ, Schweiger U, Weber B, Korner A, Standhardt H, Gotthardt U, Schmider J, Pflaum CD, Heuser I (1997) Insulin-like growth factor-I (IGF-I) plasma concentrations are increased in depressed patients. Psychoneuroendocrinology 22:493–503.

Fernandez AM, Torres-Aleman I (2012) The many faces of insulin-like peptide signalling in the brain. Nat Rev Neurosci 13:225–239.

Flores Á, Saravia R, Maldonado R, Berrendero F (2015) Orexins and fear: implications for the treatment of anxiety disorders. Trends in Neurosciences 38:550–559.

Harris GC, Wimmer M, Aston-Jones G (2005) A role for lateral hypothalamic orexin neurons in reward seeking. Nature 437:556–559.

Hoshaw BA, Hill TI, Crowley JJ, Malberg JE, Khawaja X, Rosenzweig-Lipson S, Schechter LE, Lucki I (2008) Antidepressant-like behavioral effects of IGF-I produced by enhanced serotonin transmission. Eur J Pharmacol 594:109–116.

Ji M-J, Zhang X-Y, Chen Z, Wang J-J, Zhu J-N (2019) Orexin prevents depressive-like behavior by promoting stress resilience. Molecular Psychiatry 24:282–293.

Johnson PL, Truitt W, Fitz SD, Minick PE, Dietrich A, Sanghani S, Traskman-Bendz L, Goddard AW, Brundin L, Shekhar A (2010) A key role for orexin in panic anxiety. Nat Med 16:111–115.

Kim YK, Na KS, Hwang JA, Yoon HK, Lee HJ, Hahn SW, Lee BH, Jung HY (2013) High insulin-like growth factor-1 in patients with bipolar I disorder: A trait marker? J Affect Disord.

Kondo M, Koyama Y, Nakamura Y, Shimada S (2017) A novel 5HT3 receptor-IGF1 mechanism distinct from SSRI-induced antidepressant effects. Mol Psychiatry.

Lin F, Suhr J, Diebold S, Heffner KL (2014) Associations between depressive symptoms and memory deficits vary as a function of insulin-like growth factor (IGF-1) levels in healthy older adults. Psychoneuroendocrinology 42:118–123.

Mahler SV, Moorman DE, Smith RJ, James MH, Aston-Jones G (2014) Motivational activation: a unifying hypothesis of orexin/hypocretin function. Nature neuroscience 17:1298–1303.

Matsuki T, Nomiyama M, Takahira H, Hirashima N, Kunita S, Takahashi S, Yagami K, Kilduff TS, Bettler B, Yanagisawa M, Sakurai T (2009) Selective loss of GABA(B) receptors in orexin-producing neurons results in disrupted sleep/wakefulness architecture. Proc Natl Acad Sci U S A 106:4459–4464.

Pfaffl MW (2001) A new mathematical model for relative quantification in real-time RT-PCR. Nucleic Acids Res 29:e45.

Pietrzak RH, el-Gabalawy R, Tsai J, Sareen J, Neumeister A, Southwick SM (2014) Typologies of posttraumatic stress disorder in the U.S. adult population. Journal of Affective Disorders 162:102–106.

Sakurai T (2014) The role of orexin in motivated behaviours. Nat Rev Neurosci 15:719–731.

Santi A, Genis L, Torres Aleman I (2018a) A Coordinated Action of Blood-Borne and Brain Insulin-Like Growth Factor I in the Response to Traumatic Brain Injury. Cereb Cortex 28:2007–2014.

Santi A, Bot M, Aleman A, Penninx BWJH, Aleman IT (2018b) Circulating insulin-like growth factor I modulates mood and is a biomarker of vulnerability to stress: from mouse to man. Translational Psychiatry 8:142.

Shalev A, Liberzon I, Marmar C (2017) Post-Traumatic Stress Disorder. New England Journal of Medicine 376:2459–2469.

Soya S, Sakurai T (2018) Orexin as a modulator of fear-related behavior: Hypothalamic control of noradrenaline circuit. Brain Res.

Soya S, Takahashi TM, McHugh TJ, Maejima T, Herlitze S, Abe M, Sakimura K, Sakurai T (2017) Orexin modulates behavioral fear expression through the locus coeruleus. Nat Commun 8:1606.

Ulrich-Lai YM, Herman JP (2009) Neural regulation of endocrine and autonomic stress responses. Nature Reviews Neuroscience 10:397–409.

van Varsseveld NC, van Bunderen CC, Sohl E, Comijs HC, Penninx BW, Lips P, Drent ML (2015) Serum insulin-like growth factor 1 and late-life depression: a population-based study. Psychoneuroendocrinology 54:31–40.

Yamanaka A, Beuckmann CT, Willie JT, Hara J, Tsujino N, Mieda M, Tominaga M, Yagami K, Sugiyama F, Goto K, Yanagisawa M, Sakurai T (2003) Hypothalamic orexin neurons regulate arousal according to energy balance in mice. Neuron 38:701–713.

Yurgil KA, Barkauskas DA, Vasterling JJ, Nievergelt CM, Larson GE, Schork NJ, Litz BT, Nash WP, Baker DG (2014) Association between traumatic brain injury and risk of posttraumatic stress disorder in active-duty Marines. JAMA Psychiatry 71:149–157.

