## Supplementary figures and images for "Insulin-Like Growth Factor I Modulates Vulnerability to Stress Through Orexin Neurons"

### Supplementary Figure

#### Slide 1
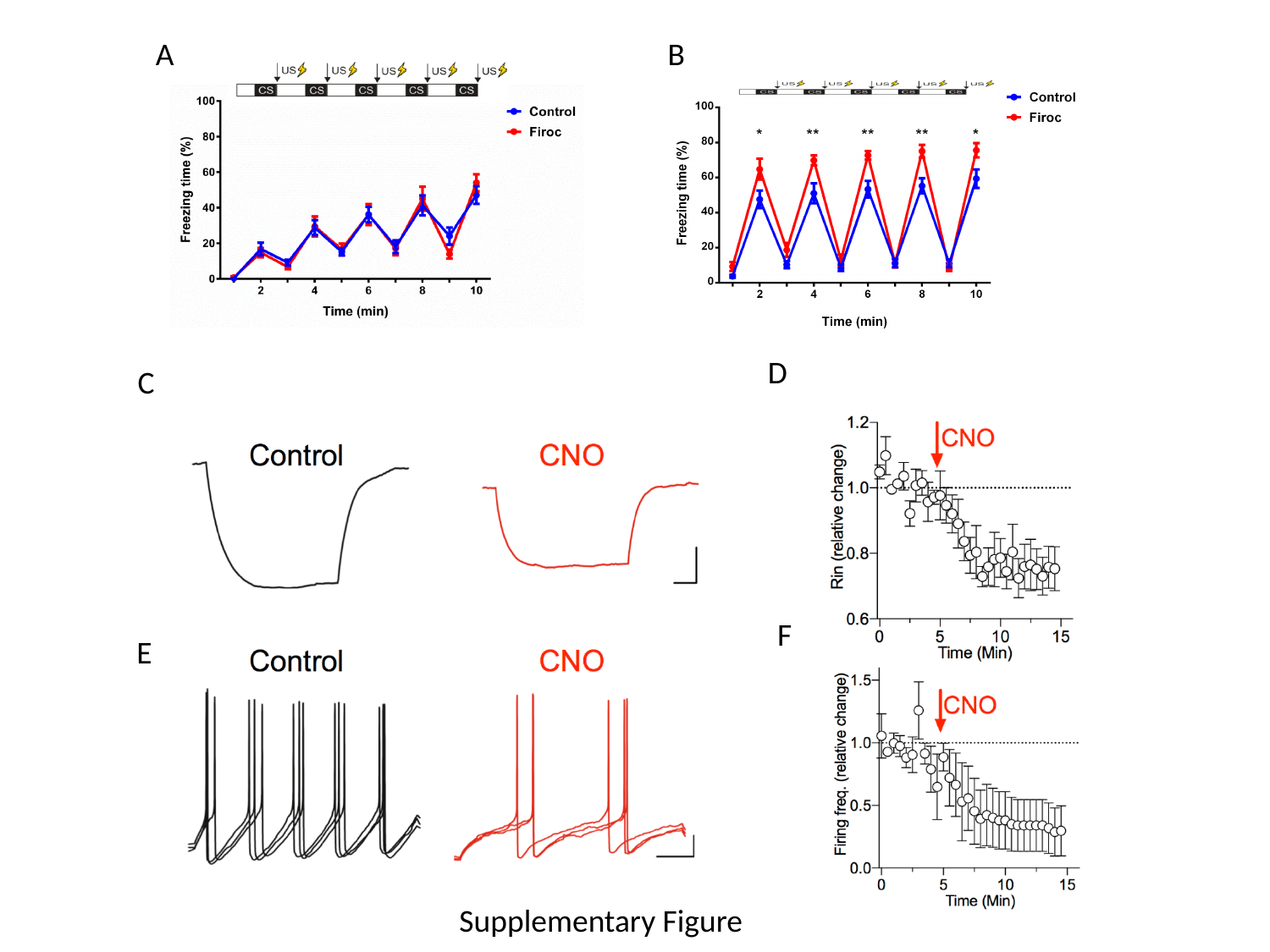

A
B
D
C
F
E
Supplementary Figure
